# Analysis of gene expression and connectivity on hippocampus of Alzheimer’s disease by a new comprehensive approach

**DOI:** 10.1101/2020.01.14.906446

**Authors:** Shan Jiang

## Abstract

Gene expression and gene connectivity describe two different functional aspects of a gene. These two different measures reveal different information about the involvement of genes in disorders. Previous case-control gene expression studies have often focused on expression level of individual genes. Correlated expression relationships among genes, measured as gene connectivity, have obtained limited attention. We developed a comprehensive method, TRIple Differentiation (TRID), to assess these two measures, both separately and jointly. We applied TRID to gene expression data in hippocampus tissue samples from three Alzheimer’s disease (AD) microarray datasets. Following TRID, comparisons among the three datasets showed poor consistency for disease-associated individual genes but reproducible changes of disease-associated biological pathways annotated for functional protein-protein interaction (PPI) modules identified from network analysis. Our results suggest that changes of gene expression in hippocampus of AD patients are highly heterogeneous at the individual gene level, while biological pathways annotated for PPI modules identified based on TRID weights demonstrate consistency among the three datasets. The R package TRID can be accessed from GitHub (https://github.com/shannjiang/TRID).

## 1. Introduction

Investigating change of gene expression level (inner trait of a gene) in patients in contrast to healthy controls is the conventional approach to identify disease-associated gene. However, differential expression does not capture change with respect to gene relationship (inter trait of a gene) between cases and controls. To study change of gene relationship, Horvath and colleagues developed Weighted Gene Co-expression Network Analysis (WGCNA) a decade ago. In common practice, WGCNA identifies clusters of co-expressed genes based on correlated expression. Differential expression between cases and controls is then evaluated for eigengene of each module, which represent the expression level of an entire module (Langfelder and Horvath, 2008; Zhang and Horvath, 2005). Therefore, WGCNA still examines change of expression level in nature; the concept of gene relationship only applied to the construction of co-expression network.

To supplement WGCNA, Fuller *et al.* introduced the concept and calculation of gene connectivity (Fuller et al., 2007). Gene connectivity was defined as the summed co-expression levels of a specific gene with all other genes (Fuller et al., 2007). Gene connectivity can be calculated in cases and controls separately and then compared to assess differential connectivity of individual genes (Fuller et al., 2007). Differential connectivity has been applied successfully (Doering et al., 2012; Fuller et al., 2007).

However, analyzing either differential expression or differential connectivity alone may miss genes whose changes are not significant uni-dimensionally but stand out if and when assessed jointly. It is desirable to investigate disturbance of genes with regards to differential expression and differential connectivity simultaneously to avoid the missed opportunity.

In this study, we developed a new method TRIple Differentiation (TRID), which includes three analyses: differential expression, differential connectivity and a fusion analysis of both differential expression and differential connectivity. The fusion analysis was termed Differential COnnectivity and Differential Expression (DiCODE). We comprehensively assessed changes of genes in hippocampus of Alzheimer’s disease (AD) patients and identified highly consistent biological processes or pathways among multiple datasets by TRID.

## 2. Methods

### 2.1. AD hippocampus expression data collection and quality control

Microarray expression data regarding hippocampus of AD patients were retrieved from Gene Expression Omnibus (http://www.ncbi.nlm.nih.gov/geo) by searching with the keywords “Alzheimer” and “hippocampus”. To reduce uncontrollable bias, only microarray datasets with consistent assay platform were included. After manual inspection and filtration, three datasets containing hippocampus expression data, GSE5281 (10 AD cases and 13 controls), GSE28146 (22 AD cases and 8 controls) and GSE48350 (19 AD cases and 43 controls), were retained for analysis.

Robust multichip average (RMA) normalization was performed on the three microarray datasets using R package affy (version 1.57.0).

### 2.2. Gene weighting strategies

To apply TRID to the three datasets, we began with the classical differential expression analysis between cases and controls, which was termed as the differential expression test in TRID. Welch’s t-test was used to assess differential expression levels of individual genes between cases and controls (Doering et al., 2012). Absolute value of the t-statistic, denoted by |*T*(*i*)|, was used to represent the differential expression (DE) weight of the *i*th gene, where a higher absolute t-statistic value corresponds to a greater differential expression level.

For the differential connectivity (DC) test in TRID, connectivity levels of individual genes in cases and controls were calculated respectively by using 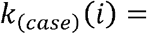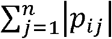 and 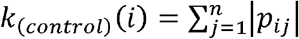, where the *p*_*ij*_ denotes the Pearson correlation coefficient between the *i*th and *j*th genes. To facilitate the comparison of connectivity level of the *i*th gene between cases and controls, connectivity level of the *i*th gene in cases and controls were standardized through z-score transformation such that 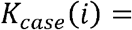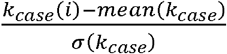 and 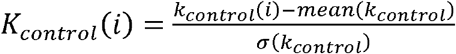. *DiffK*(*i*) = *K*_*case*_(*i*) – *K*_*control*_(*i*) was defined as differential connectivity of the *i*th gene and |*Dif fK*(*i*)| as the differential connectivity weight of the *i*th gene, as defined by Fuller *et al* (Fuller et al., 2007).

For the combined Differential COnnectivity and Differential Expression (DiCODE) test in TRID, in order to fuse differential expression and differential connectivity weights calculated for the *i*th gene, the differential expression weight |*T*(*i*)| and differential connectivity weight |*DiffK*(*i*)| were standardized through z-score transformation such that 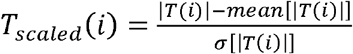 and 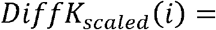 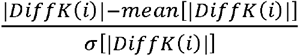. *DiCODE*(*i*) = *T*_*scaled*_(*i*) + *DiffK*_*scaled*_(*i*) was defined as the DiCODE weight of the *i*th gene.

To evaluate the statistical significance of the gene weight individually, for a weighting strategy, we randomly sampled the gene weight for 1 million times to generate a reference distribution. Gene weight *P*-value was calculated as the proportion of sampled gene weights as or more extreme than the given gene weight.

The R package TRID can be downloaded from GitHub (https://github.com/shannjiang/TRID).

### 2.3. TRID weight-based functional PPI module identification and gene ontology (GO) annotation

DE, DC and DiCODE weights were termed TRID weights in this study. Cytoscape plugin jActiveModules was used to identify significantly enriched functional PPI modules based on TRID weights of genes (Ideker et al., 2002). In this study, we used PPI information from the IntAct and MINT databases, which detail molecular interactions in well-studied organisms such as *S. cerevisiae* and *Homo sapiens*, to construct a combined background PPI network (Cline et al., 2007; Hermjakob et al., 2004; Zanzoni et al., 2002).

DAVID was then used to annotate the functions of identified PPI modules (Huang et al., 2009).

## 3. Results

### 3.1. TRID weights of each gene generated from hippocampus microarray datasets

For TRID weights of individual genes and their respective *P*-values generated from GSE5281, GSE28146 and GSE48350 microarray datasets, please refer to supplementary Table 1, 2 and 3 respectively.

For DE, DC and DiCODE weighting strategies, venn diagrams of shared and specific top 5% high ranking weighted genes among the three datasets were shown in Fig. 1. For DE weighting strategy, 19 genes were shared among the three datasets (Fig. 1A). For both DC and DiCODE weighting strategy, only 3 genes were shared among the three datasets (Fig. 1B and 1C).

**Fig. 1.**
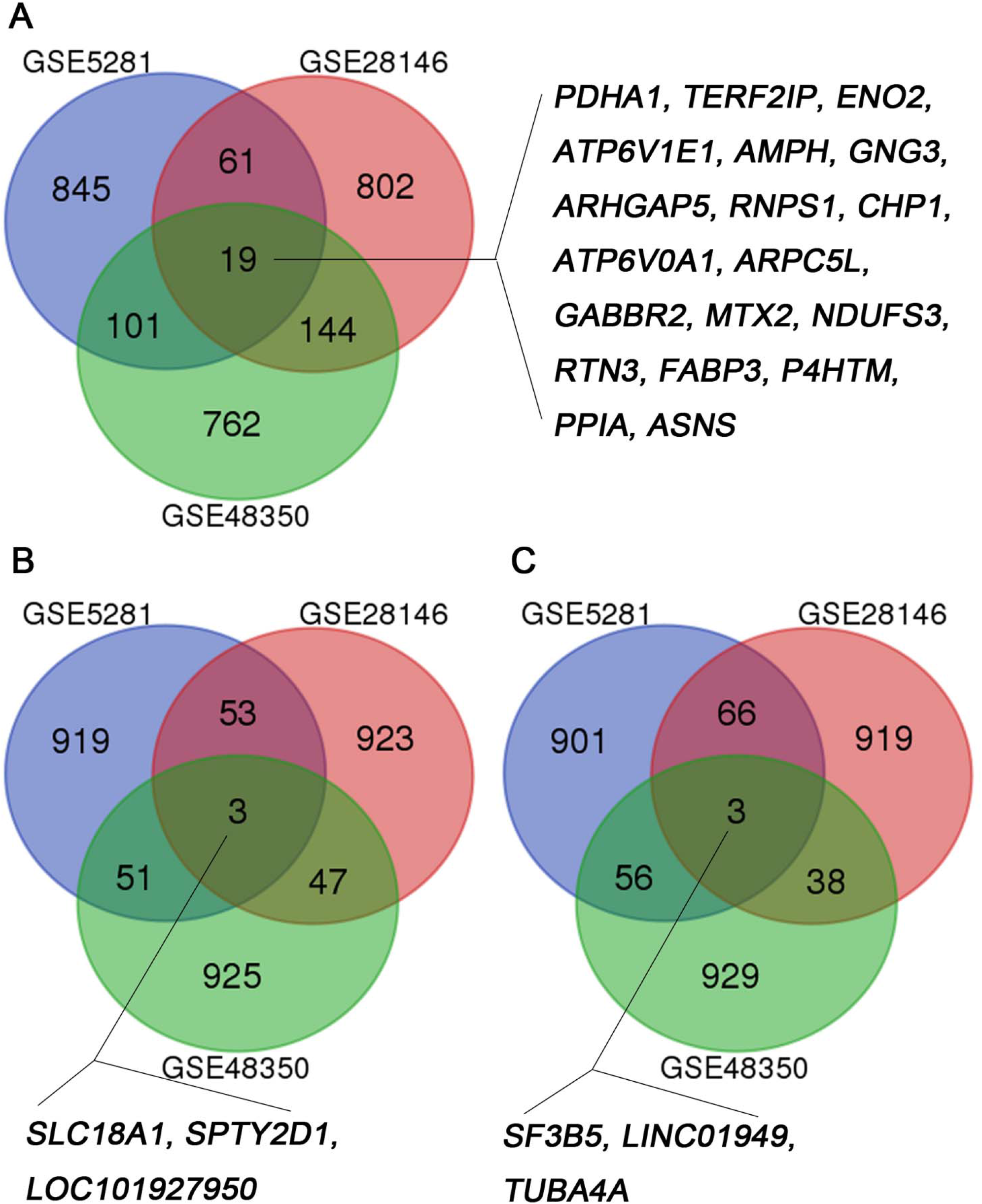
Venn diagrams of shared and specific top 5% high ranking weighted genes among the three hippocampus microarray datasets based on TRID DE (A), DC (B) and DiCODE (C) weighting strategies.

### 3.2. TRID weights-based functional PPI module identification and GO annotation

TRID weights-based functional PPI modules were identified for individual datasets. For the network graphs of functional PPI modules of individual datasets identified based on DE, DC and DiCODE weights, please refer to Supplementary Figures 1, 2, and 3 respectively. For detailed GO annotations of these identified functional PPI modules of individual datasets based on DE, DC and DiCODE weights, please refer to Supplementary Table 4, 5 and 6 respectively. Venn diagrams of shared and specific significantly enriched GO terms among the three datasets based on DE, DC and DiCODE weights were shown in Fig. 2A, 2B and 2C respectively. Nearly 30 to 50% of the significantly enriched GO terms were shared among the three datasets based on each weight of TRID. For detailed significant GO terms shared among the three datasets please refer to Supplementary Tables 7, 8 and 9 for DE, DC and DiCODE weighting strategies respectively. Cell nucleus-associated biological processes, cell components and molecular functions were highly shared for individual weighting strategies of TRID. Nuclear lamins are fibrous proteins providing structural function and transcriptional regulation in the cell nucleus (Dechat et al., 2008). Dysfunctions of and mutations in the genes encoding A or B-type lamins, or termed laminopathy, were associated to neurodegeneration recently (Frost, 2016; Frost et al., 2016).

**Fig. 2.**
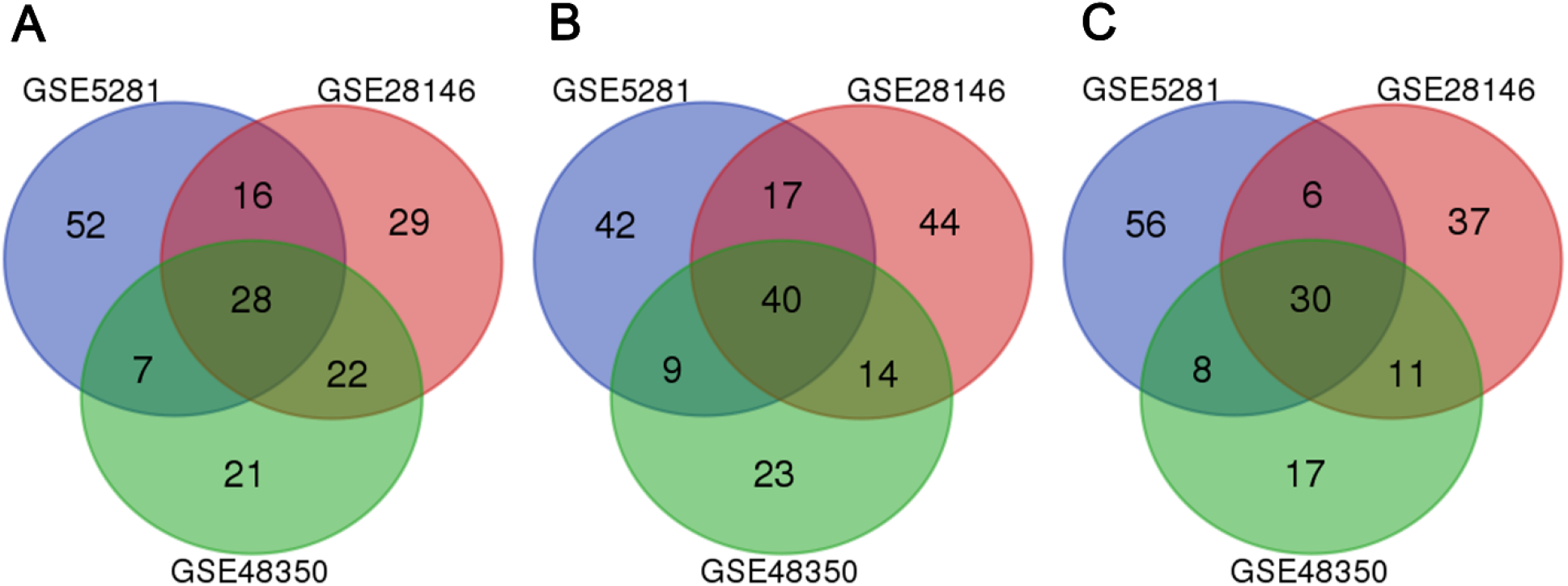
Venn diagrams of shared and specific GO term annotations for identified functional PPI modules based on TRID DE (A), DC (B) and DiCODE (C) weights.

## 4. Discussion

In this study we developed a new method called TRID, which includes differential expression analysis, differential connectivity analysis and DiCODE analysis for individual genes. With DiCODE analysis, we can detect genes with changes that could be missed in conventional differential expression analysis, emphasizing the combined effects instead of either one alone.

Only less than 2% of top 5% high TRID-weighted genes were shared among the three datasets (Fig. 1), which suggests heterogeneity among samples. More than 90% of disease-associated SNPs are located in non-coding regions of the genome (Hindorff et al., 2009), suggesting the potential regulatory nature of these SNPs. The International Genomics of Alzheimer’s Project (IGAP) GWAS meta-analysis identified 20 loci associated with AD (Lambert et al., 2013). In this study, most of the AD hits lie outside of gene coding regions. At least some of them may contribute to changes in gene expression (Knight, 2003). Variable combinations of risk alleles at different non-coding SNP loci in patients contribute to expression profile heterogeneity in AD. Moreover, environment, including social and psychological factors (Read et al., 2004), can potentially influence gene expression. Some research have implied that epigenetic modifications largely influence regulations of genes in AD (Kwok, 2010; Rao et al., 2012), which could further contribute to AD heterogeneity or individual differences of expression profiles.

In contrast to the high heterogeneity at individual gene level, we observed very robust findings of biological processes or signaling pathways from functional PPI module identification and annotation, even though all datasets used in this study had relatively small sample sizes. It indicates that different genes responsible for the underlying etiology of AD involve in the same pathways or PPI modules. Methodologically, by coupling to TRID weighting, functional PPI module identification and annotation could be an effective way to uncover pathways involved in AD progression.

In summary, our study revealed that (1) TRID can weight genes effectively and comprehensively; (2) coupling PPI module analyses with TRID weighting may help to detect the converged changes in AD; and (3) AD is heterogeneous at individual gene level, but many differential genes are involved in the same nucleus-associated pathways and disease regulatory modules.

## Conflict of interest

The authors declare no conflict of interest.

## Notes

### Competing Interest Statement

The authors have declared no competing interest.

### Summary of Updates

Corresponding address

