## Supplementary Figures for "Analysis of gene expression and connectivity on hippocampus of Alzheimer’s disease by a new comprehensive approach"

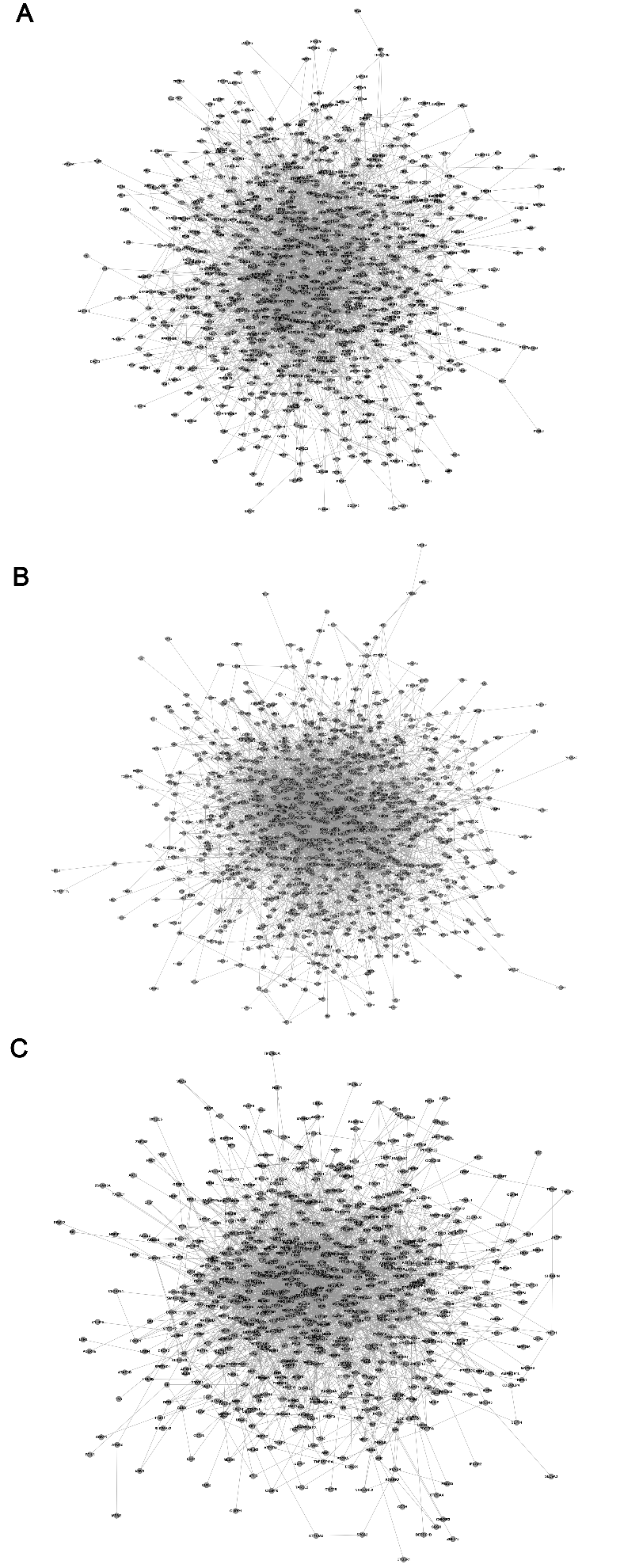

**Supplementary Figure 1.** Network graphs of functional PPI modules identified based on DE weight of TRID using GSE5281 (A), GSE28146 (B) and GSE48350 (C) datasets respectively.

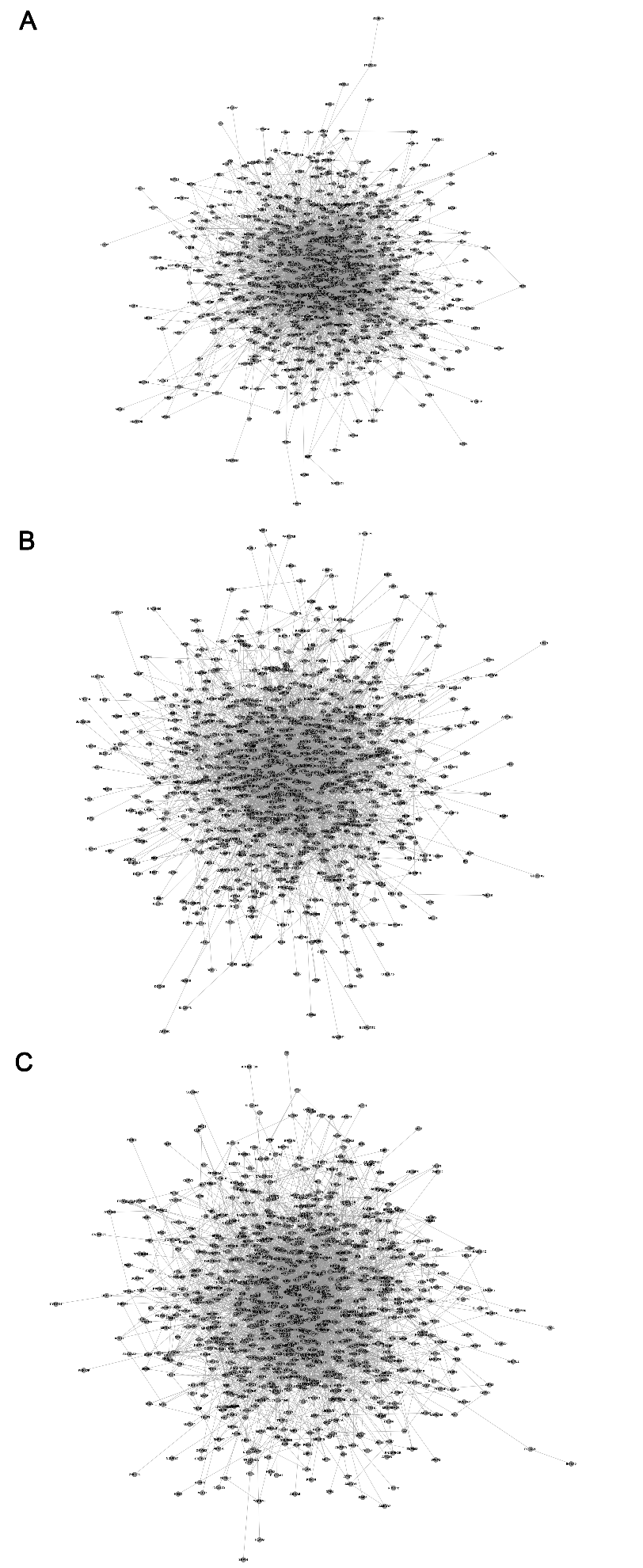

**Supplementary Figure 2.** Network graphs of functional PPI modules identified based on DC weight of TRID using GSE5281 (A), GSE28146 (B) and GSE48350 (C) datasets respectively.

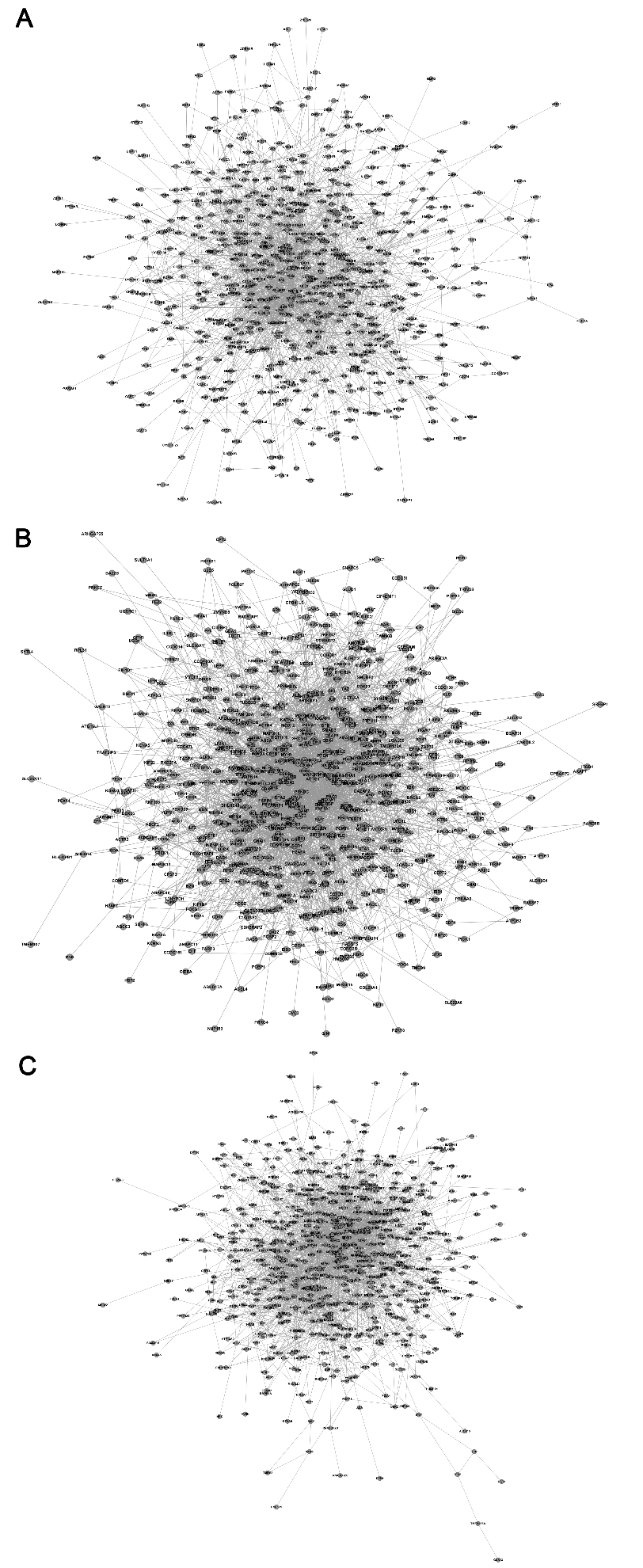

**Supplementary Figure 3.** Network graphs of functional PPI modules identified based on DiCODE weight of TRID using GSE5281 (A), GSE28146 (B) and GSE48350 (C) datasets respectively.
