## Supplementary Table 4-6 legends for "Analysis of gene expression and connectivity on hippocampus of Alzheimer’s disease by a new comprehensive approach"

Supplementary Table 4. Full lists of GO annotations of identified functional PPI modules based on DE weight for individual datasets

Supplementary Table 5. Full lists of GO annotations of identified functional PPI modules based on DC weight for individual datasets

Supplementary Table 6. Full lists of GO annotations of identified functional PPI modules based on DiCODE weight for individual datasets
